# Efficient differentiation and polarization of primary cultured neurons on poly(lactic acid) scaffolds with microgrooved structures

**DOI:** 10.1101/644781

**Authors:** Asako Otomo, Mahoko Takahashi Ueda, Toshinori Fujie, Arihiro Hasebe, Yosuke Okamura, Shinji Takeoka, Shinji Hadano, So Nakagawa

## Abstract

Synthetic biodegradable polymers including poly(lactic acid) (PLA) are attractive cell culture substrates because their surfaces can be micropatterned to support cell adhesion. The cell adhesion properties of a scaffold mainly depend on its surface chemical and structural features; however, it remains unclear how these characteristics affect the growth and differentiation of cultured cells or their gene expression. In this study, we fabricated two differently structured PLA nanosheets: flat and microgrooved. We assessed the growth and differentiation of mouse primary cultured cortical neurons on these two types of nanosheets after pre-coating with poly-D-lysine and vitronectin. Interestingly, prominent neurite bundles were formed along the grooves on the microgrooved nanosheets, whereas thin and randomly extended neurites were only observed on the flat nanosheets. Comparative RNA sequencing analyses revealed that the expression of genes related to postsynaptic density, dendritic shafts, and asymmetric synapses was significantly and consistently up-regulated in cells cultured on the microgrooved nanosheets when compared with those cultured on the flat nanosheets. These results indicate that microgrooved PLA nanosheets can provide a powerful means of establishing a culture system for the efficient and reproducible differentiation of neurons, which will facilitate future investigations of the molecular mechanisms underlying the pathogenesis of neurological disorders.

## Introduction

Dissociated primary neuronal cultures are widely used not only for basic neuroscience research but also for drug discovery for neurological disorders^1,2,3^. In such culture systems, scaffolds are one of the key factors providing the cells with structural support for attachment and subsequent growth and differentiation. Thus far, numerous synthetic polymers including polystyrene, poly(lactic acid) (PLA), poly(glycolic acid), and poly(lactic-co-glycolic acid)^4,5,6^ have been developed to serve as scaffolds. Among these, PLA, a biodegradable and resorbable polyester, has recently come into the limelight for its utility in medical applications such as tissue regeneration^7^.

Polymeric ultrathin film consisting of PLA, hereinafter called “PLA nanosheet,” is a thin, soft, and flexible material, with properties that allow it to adhere anywhere without any adhesive materials.^8^ Many studies have demonstrated that nanosheets can be used to dress wounds to avoid suture, prevent infection, and bone regeneration etc. for biomedical applications^8,9,10,11,12,13^. Nanosheets are also suitable for use as a sheet substrate in cell culture for several reasons. First, nanosheets can easily adhere to the surface of standard culture plates, culture dishes, and cover glass without any adhesive materials. Second, cells and/or tissues cultured on nanosheets can be easily recovered, allowing researchers to easily analyze biological molecules such as proteins, DNA, and RNA. Last, a variety of structural patterns of the nanosheet surface, such as grooves and pores, is possible^14,15^.

Despite these advantages, there are a number of issues that can hinder the application of nanosheets to cell culture experiments. The surface of the nanosheet is hydrophobic, which prevents cell adhesion. Therefore, surface pre-treatment of the nanosheets is required^16,17,18^. For cells, particularly dissociated neurons, cell adhesion molecules such as poly-D-lysine (PDL) peptides, which confer a positive charge on the nanosheet surface and assist cell adhesion^19^, are required. Further, dissociated cultured neurons under standard culture conditions^20^ extend neurites in random directions, which prevents neurons from forming organized neuronal networks. This may mainly be due to a lack of appropriate attractive and repulsive biological cues from the surrounding cells as well as an absence of scaffold-linked mechanical cues to guide the direction of axon pathfinding. Moreover, it has been shown that morphogenesis in cultured neurons can be affected by topographical differences on the PLA substrate, and grooved structures, in particular, may improve the guidance of neurite extension^21^, although the molecular mechanisms underlying such phenomena remain to be investigated.

In this study, we fabricated microgrooved nanosheets with different microgroove widths and used flat nanosheets as the control. After coating with cell adhesion molecules, we then investigated the effects of the topographical features of these nanosheets on the morphology of mouse primary cultured cortical neurons. Further, to elucidate the molecular basis of the observed differences, we compared the gene expression of cell cultures on the two types of nanosheets. Our findings indicated that microgrooved nanosheets serve as an effective scaffold for the controlled neurite polarization of cultured neurons, thereby promoting the efficient and reproducible differentiation of neurons. Thus, microgrooved nanosheets are expected to be applied to a large number of investigations in neuroscience research as well as regenerative medicine.

## Results and Discussions

### Assessment of materials used to pre-coat the nanosheet surface

The surface of a PLA nanosheet is smooth and hydrophobic, thereby preventing cell adhesion^16,18^. To establish the appropriate conditions for pre-coating the surface of the nanosheets, we assessed PDL and PDL coated over truncated recombinant human vitronectin (PDL + VTN-N). VTN-N is a recombinant human protein that can provide a defined surface for feeder-free culture of human pluripotent stem cells^22^. VTN-N is an extracellular matrix molecule that supports neurite outgrowth *in vitro* both under normal conditions and after trauma^23^. We then cultured PC12 cells, a cell line derived from pheochromocytoma in the rat adrenal medulla, on a glass coverslip or nanosheet pre-coated with PDL or PDL + VTN-N. PC12 cells attached and grew on the glass as well as the nanosheets that were pre-coated with either PDL or PDL + VTN-N (Figure 1A). In contrast, without any PDL coating, the PC12 cells hardly adhered to the nanosheet (Figure 1A). We further confirmed that mouse primary cultured cortical neurons grew and differentiated on the nanosheets coated with PDL or PDL + VTN-N as was observed for the PC12 cells (Figure 1B). At 2 days *in vitro* (DIV2), cell adhesion and neurite protrusion were observed for mouse primary cortical neurons on the nanosheet coated with either PDL or PDL + VTN-N (Figure 1B). Notably, PDL + VTN-N coating particularly enhanced cell adhesion of the primary cultured cortical neurons on the nanosheet (Figure 1B). At 6 days *in vitro* (DIV6), this observation became even more evident: the degree of neurite extension on the PDL + VTN-N-coated nanosheet was much higher than that on the PDL-coated one (Figure 1B). At this time point, neuron-extended neurites were beginning to connect to each other (Figure 1B), suggesting that normal differentiation of the neurons was achieved on the nanosheet. Moreover, the cell viability of the cultured cortical neurons on the nanosheet at DIV6 reached approximately 92%, which was comparable with that on a conventional glass substrate (Figure 1C). Taken together, the nanosheet coated with PDL + VTN-N is likely suitable for neuronal cultures.

**Figure 1.**
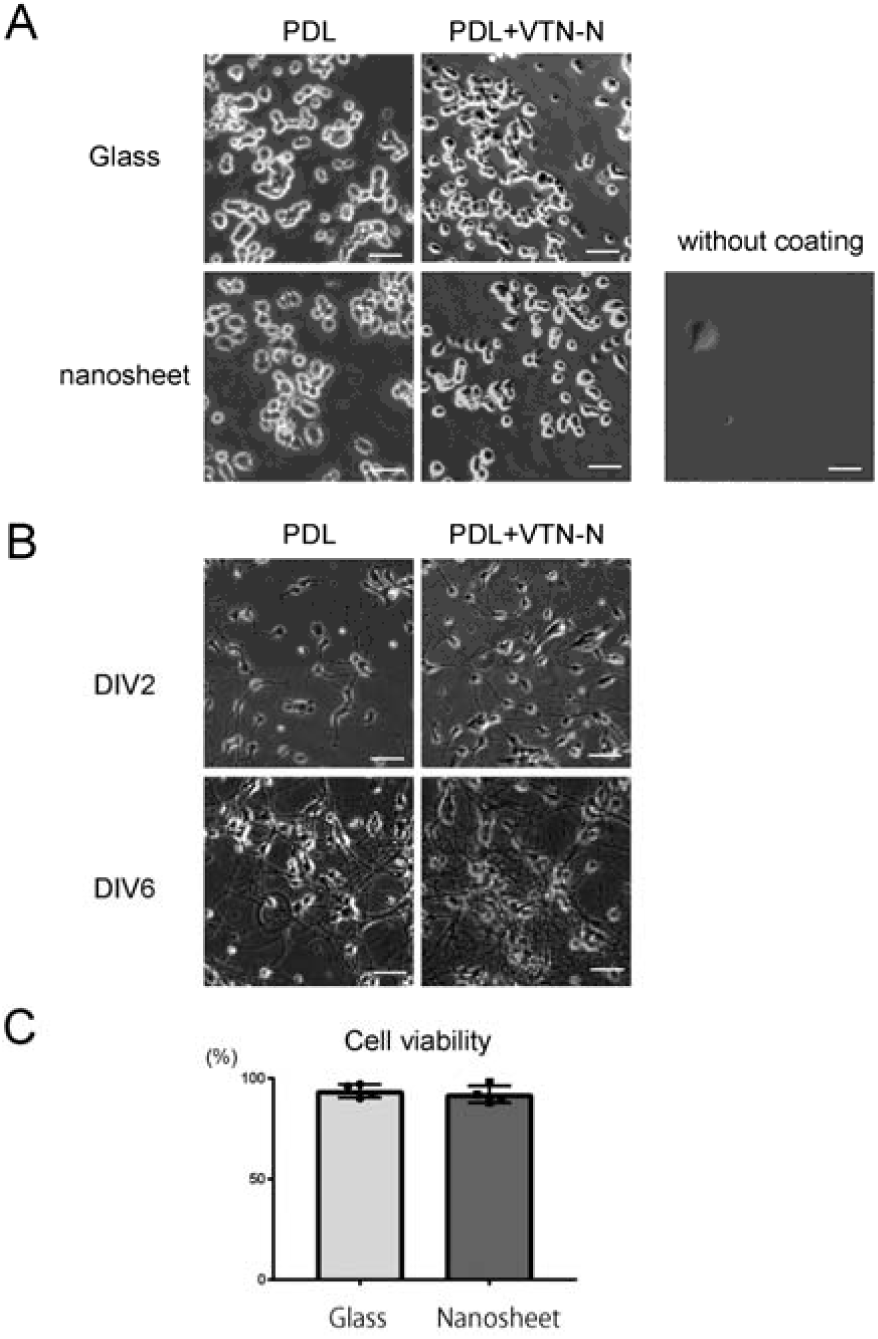
Culturing neurons on the PLA nanosheet (Figure legend up to 350 Words) A) PC12 cells cultured on a glass substrate or PLA nanosheet. The glass substrate and PLA nanosheet were coated with PDL or PDL + VTN-N. In the absence of any coating, the PC12 cells hardly adhered to the PLA nanosheet. Scale bars, 50 *µ*m. B) Mouse primary cultured cortical neurons on a PDL + VTN-N-coated PLA nanosheet. Neural morphology at DIV2 and DIV6 are shown. Mouse primary cultured cortical neurons adhered to and displayed more prominently elongated neurites on the PDL + VTN-N-coated PLA nanosheet. Scale bars, 50 *µ*m. C) Cell viability of mouse primary cultured neurons on the glass substrate and PLA nanosheet were compared with neurons cultured on conventional plastic culture dishes coated with PDL + VTN-N. The cell viability of neurons on the PLA nanosheet was comparable with that on the glass substrate [glass: 93.75 ± 1.65, nanosheet: 92.00 ± 2.16 (mean ± SE), expressed as a percentage of the cell viability on plastic culture dishes].

### Development of the microgrooved nanosheet

To develop the nanosheet with a microgrooved surface that was applicable to neuronal cultures, we first sought to fabricate nanosheets with three different surface structures, of which the parallel microgrooves were 20, 30, or 50 μm wide with a height of 6 μm, according to previously reported procedures^24,25,26^ (Figure S1). In brief, PLA nanosheets were prepared by spin coating of the PLA-dichloromethane solution on a polydimethylsiloxane (PDMS) negative replica with microgrooved motifs. The micropatterned PLA nanosheets were overlaid with a poly(vinyl alcohol) (PVA) supporting layer and then released from the PDMS mold. The nanosheets with the PVA supporting layer were immersed into phosphate-buffered saline without Mg^2+^ or Ca^2+^ [PBS(–)] and then captured by a glass substrate.

Next, to assess the quality of the processed nanosheets, we measured the fine structure of each surface with a Dektak stylus profiler (Bruker, Billerica, MA, USA; Figure S2). It was revealed that nanosheets with 50 μm wide parallel microgrooves were most stably and reproducibly fabricated, whereas those with 20 or 30 μm in width were not (Figure S2). Therefore, we decided to use nanosheets with 50 μm wide parallel microgrooves for subsequent experiments.

### Morphological analysis of neuronal cells cultured on microgrooved nanosheets

To assess the effects of the surface microstructure of the nanosheet on the cell adhesion, neurite outgrowth, and morphology of the cultured neurons, we cultured mouse primary cortical neurons on nanosheets with either a flat or parallel-microgrooved surface that were pre-coated with PDL + VTN-N. At 9 days *in vitro*, we fixed and stained the cells with anti-MAP2 and anti-Tuj-1 antibodies and investigated their neurite orientations and morphologies on the nanosheet. Tuj-1 and MAP2 are an axon and a dendrite marker, respectively. Cultured neurons on both the flat and microgrooved nanosheets were positive for MAP2 and Tuj-1, suggesting that the cells firmly attached to and fully differentiated on the nanosheets irrespective of their surface structure (Figure 2A). Notably, cultured neurons on the microgrooved nanosheet extended neurites along the direction of the parallel microgrooves and formed thick neurite bundles (Figure 2A). In contrast, neurons on the flat nanosheet extended thin and separated neurites in random directions (Figure 2A).

**Figure 2.**
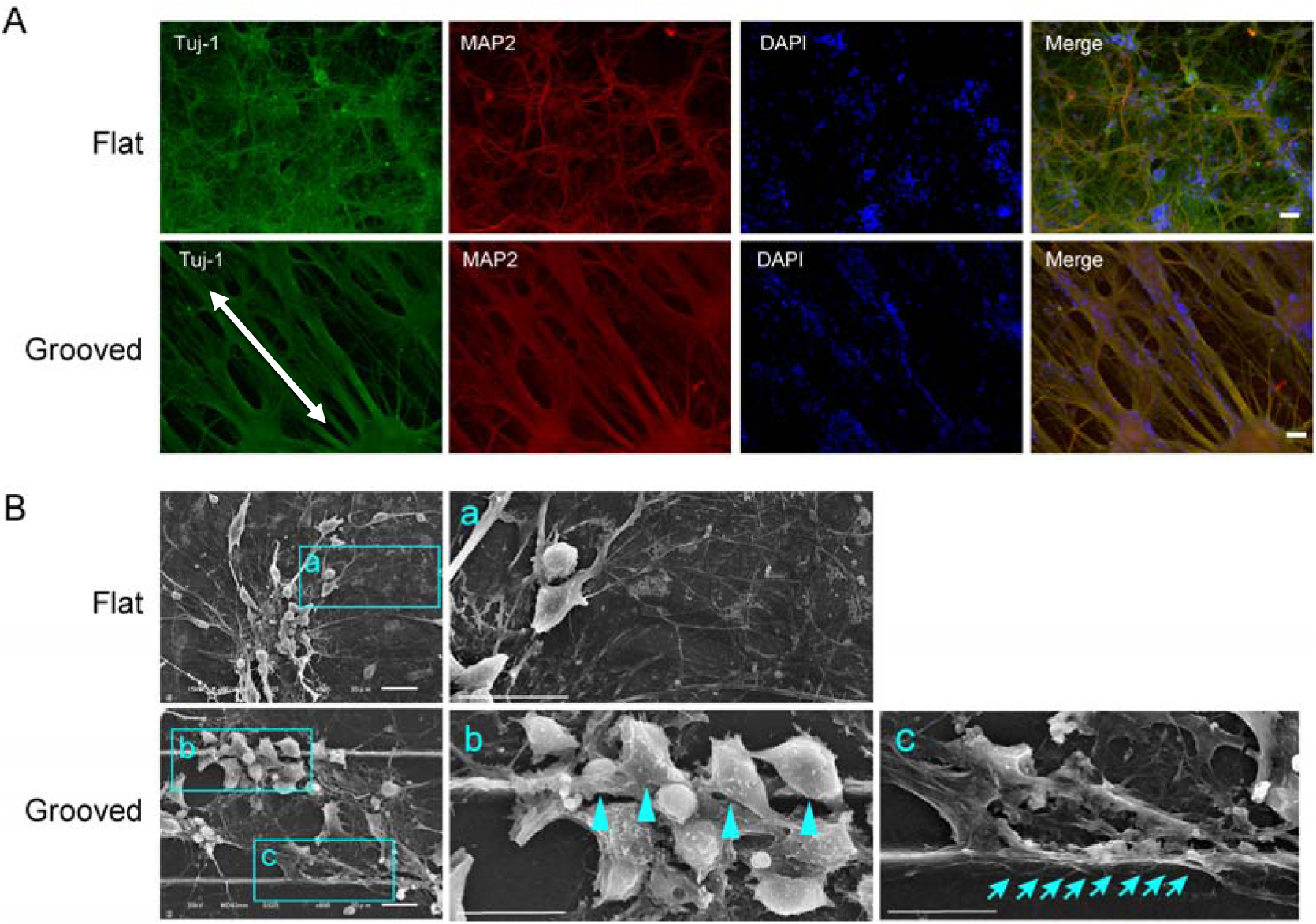
Culturing neurons on the flat and microgrooved PLA nanosheets. A) Morphological analysis of neural cell culture on nanosheets. Tuj-1 and MAP2 were used as a neurite and dendrite marker, respectively. Nuclei were visualized by DAPI staining. The white arrows indicate the direction of microgroove processing. Cultured neurons on the microgrooved nanosheet extended neurites along the microgrooves. In contrast, cultured neurons on the flat nanosheet extended neurites in random directions. Scale bars, 50 μm. B) SEM analysis of mouse primary cultured cortical neurons on flat and microgrooved PLA nanosheets. Neurons on the flat nanosheet adhered to each other and extended neurites from the cell body (a). Neurons on the microgrooved nanosheet densely colonized the inside of the microgrooves between the vertical ridges. Interestingly, neurons also adhered to the sidewall of the ridges rather than to the bottom of the microgrooves (b, indicated by blue arrowheads) and then extended neurites (c, indicated by blue arrows). Scale bars, 20 μm.

To examine how the grooved structure serves as a structural scaffold for neurons, we investigated the locations where neurons adhered to the microgroove structures by using scanning electron microscopy (SEM) (Figure 2B). For this purpose, we cultured a limited number of cells on the microgrooved nanosheet because a high-density culture (such as in Figure 2A) did not allow us to observe the location of cell adhesion. Neurons on the flat nanosheet adhered to each other and elongated neurites from the cell body in random directions (Figure 2B-a). In contrast, neurons on the microgrooved nanosheet densely colonized the inside of the microgrooves between the vertical ridges (Figure 2B). Interestingly, enlarged images revealed that some neurons appeared to adhere to the sidewall of the ridges rather than to the bottom of the microgrooves (Figure 2B-b, indicated by arrowheads) and then extended neurites (Figure 2B-c, indicated by arrows), suggesting that neurons may also use the vertical surface of the ridges as scaffolds. These results indicate that the microgrooves on a nanosheet affect the architecture of cell-to-substrate adhesion, thereby changing the neurite morphology and, presumably, also its biological function.

### Gene expression patterns of neuronal cells cultured on the nanosheets

To understand the molecular basis of the observed morphological differences in neuronal cells between the flat and microgrooved nanosheets, we performed whole exome sequencing (RNA-Seq) analyses on the primary cultured mouse cortical neurons. Seven different samples were subjected to RNA-Seq analysis: flat nanosheet, n = 3; microgrooved nanosheet, n = 3; and glass coverslip, n = 1 (Figure 3A). For each sample, 75-bp paired-end sequencing reads were mapped to the mouse reference genome (mm10), and the results are summarized in Figure 3B. To examine gene expression levels among different samples, the number of mapped reads were normalized by regularized log transformation implemented in DESeq2 program^27^.

**Figure 3.**
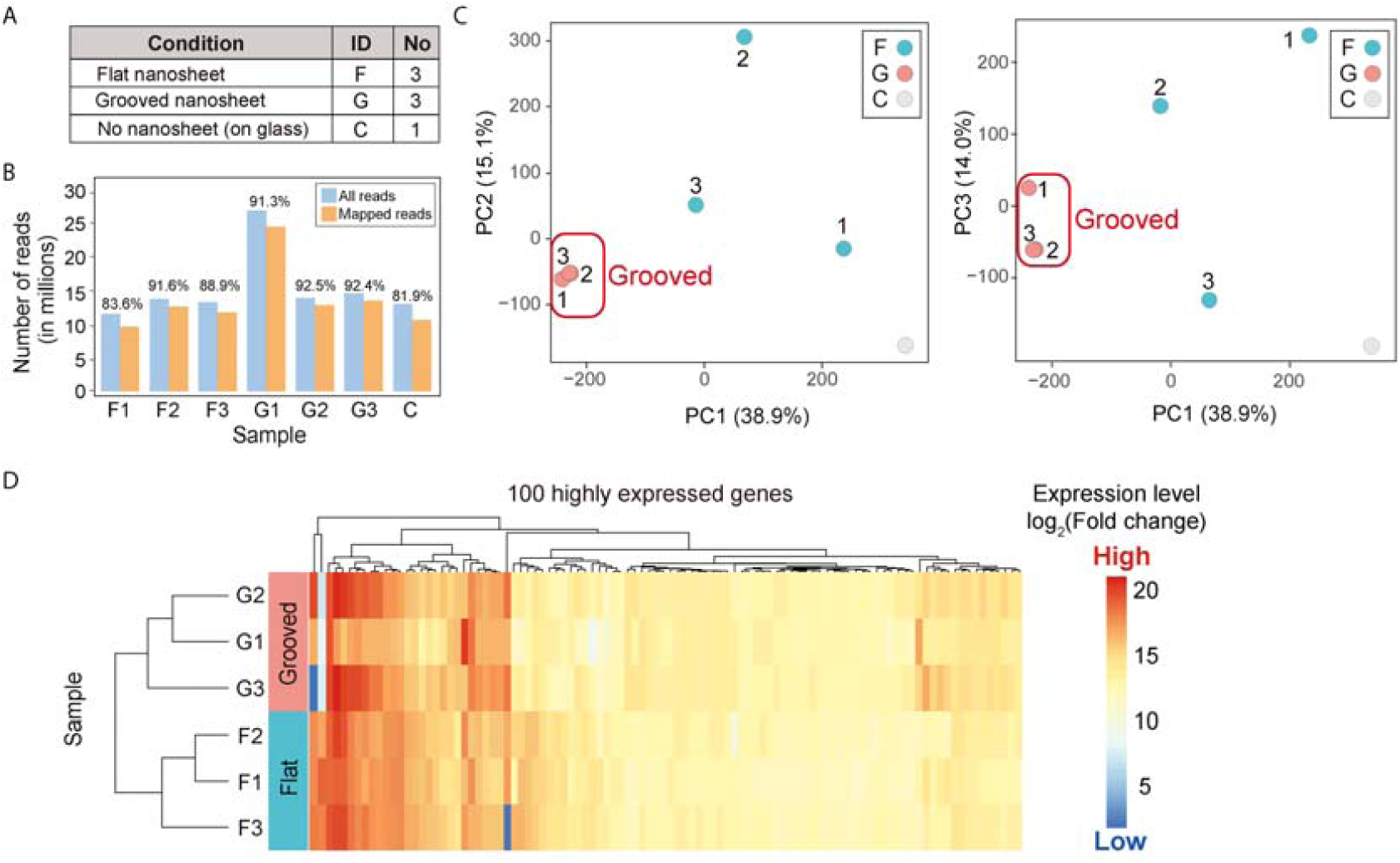
RNA-Seq analysis. A) Summary of the samples analyzed in the RNA sequencing (RNA-Seq) study. B) The number of reads per sample generated by RNA-Seq is shown. Sample names on the x-axis are the same as those in (A). A color bar indicates the number of all (blue) or mapped (orange) reads for each sample. The value above each bar represents the percentage of mapped reads. C) PCA plots of PC1, PC2, and PC3. The percentile contribution of each component’s variance is shown on each axis. The numbers above each data point indicate the numbers of the sample name. D) Heatmap of the normalized expression values of the top 100 highly expressed genes. The color indicator represents the regularized log-transformed gene expression data.

Using the normalized expression values, we assessed the overall similarity of gene expression among the seven samples by principal component analysis (PCA). PCA is a dimension-reduction method for transforming a large set of variables to small sets of principal components (PCs), which represent all the variables of a given dataset. Thus, in a PCA plot, samples forming a cluster indicate that their variations are explained by the PCs with similar patterns. PCA of the RNA-Seq data showed that the cells on the microgrooved nanosheet formed a distinct data cluster, whereas the other samples did not (Figure 3C). This indicated that neurons on the microgrooved nanosheet exhibited a consistent pattern of gene expression. The proportion of variance, which indicates how much variance was explained by each component, was 38.9%, 15.1%, and 14.0% in PC1, PC2, and PC3, respectively (Figure 3C), demonstrating that these three PCs captured approximately 68% of the gene expression variance in the samples. We further conducted clustering analysis of the normalized gene expression values of the top 100 highly expressed genes in the six nanosheet-cultured samples (Figure 3D). The results suggested that the gene expression patterns of cells on microgrooved and flat nanosheets were clearly distinct. These results combined with the morphological findings support the notion that microgrooves on the nanosheet affect the gene expression of neuronal cells and facilitate their stable differentiation.

### Differentially expressed gene analysis in neuronal cells cultured on nanosheets

To identify genes associated with the observed differences in the morphological features of neuronal cells between flat and microgrooved nanosheets, we performed differentially expressed gene (DEG) analysis. A total of 300 increased and 1,458 decreased DEGs were identified (Figure 4A). To investigate the gene functions of the DEGs potentially associated with the differences in the gene expression profiles, we performed Gene Ontology (GO) analysis. Ontology describes gene function with respect to three biological aspects: molecular function (MF), cellular component (CC), and biological process (BP) ^28^. On the basis of the GO analysis, we uncovered statistically enriched functional categories of CC and BP (Figure 4B). In particular, the functions of the up-regulated genes in samples cultured on the microgrooved nanosheet were linked to morphological features of neurons such as the postsynapse, postsynaptic density, dendritic shaft, and asymmetric synapse (Figures 4B and 4C). In contrast, the down-regulated genes were mostly linked to tubulin and microtubule binding (Figures 4B and 4C).

**Figure 4.**
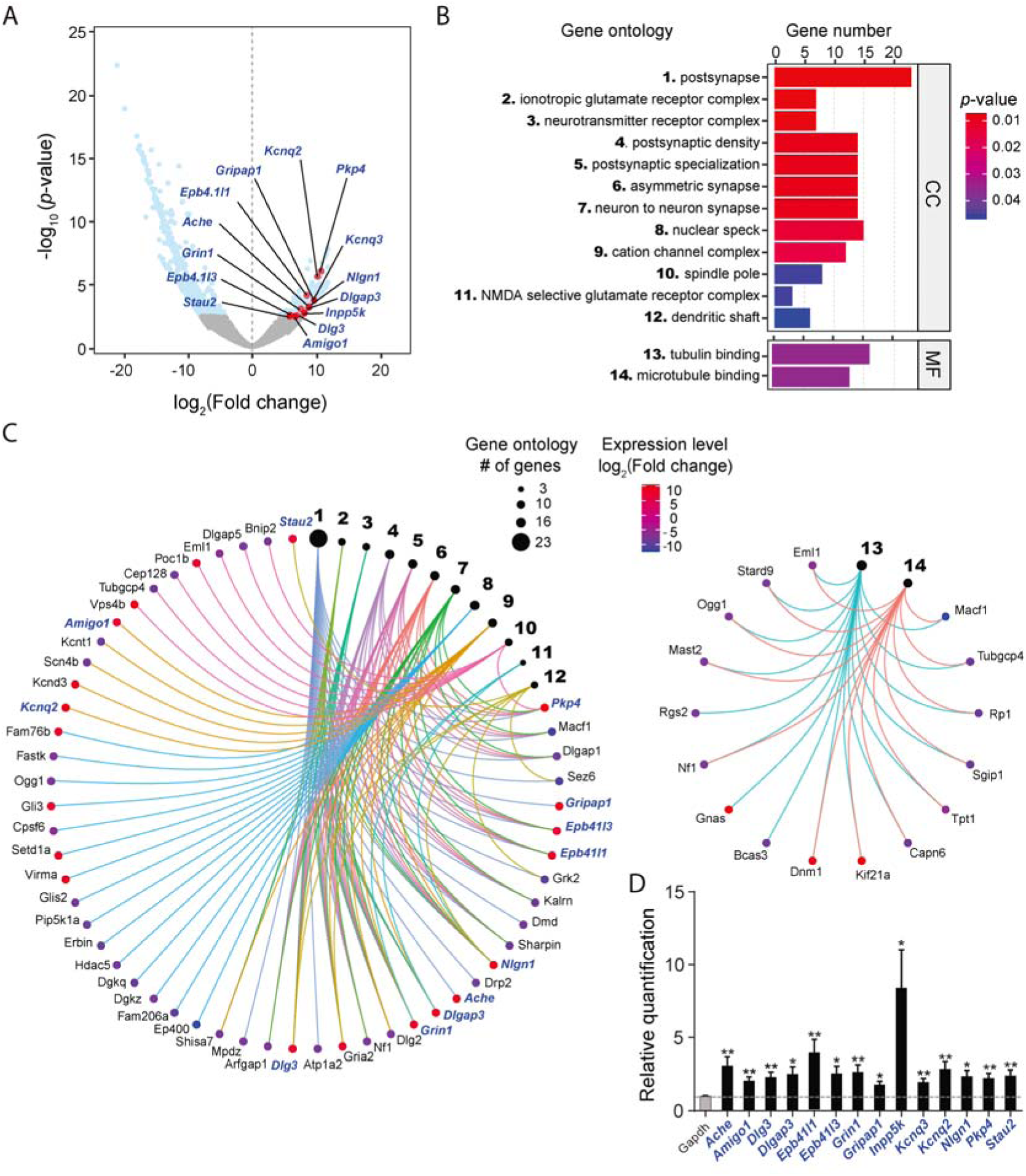
qPCR analysis of the DEGs. A) Volcano plot of the *p* value as a function of the log_2_-transformed fold changes for each gene in the samples cultured on the microgrooved versus flat nanosheets. The light blue dots represent statistically up- or down-regulated transcripts identified by DEG analysis [*p*□<□0.05, Benjamini–Hochberg (BH) corrected]. The genes validated by qPCR are highlighted in red. B) Enriched gene functions of DEG identified by Gene Ontology analysis with adjusted *p* < 0.05 (BH corrected): CC, cellular component; MF, molecular function. C) Gene-Concept Network of the CC (left) and MF (right) DEG, showing the relationship between the GO categories and the genes. The black circles represent the GO categories, whose functions are indicated by the same numbers used in (B). The size of the circles for each GO category corresponds to the number of genes in the category. The value of the log_2_-transformed fold change of each gene is indicated by the color bar. Blue italics indicate genes validated by qPCR. D) Relative gene expression levels of selected DEGs. Genes *Ache, Amigo1, Dlg3, Dlgap3, Epb41l3, Epb41l1, Grin, Gripap1, Inpp5k, Kcnd3, Kcnq2, Nlgn1, Pkp4*, and *Stau2* in the samples from the flat and microgrooved nanosheets were quantified by qPCR. The dotted line represents the gene expression of samples on the flat nanosheet. Statistically significant differences between the flat and microgrooved nanosheets are indicated by asterisks (\*\**p* < 0.01, \**p* < 0.05) (Fig. 4D). Expression of *Gapdh* was used as an internal control.

The up-regulated genes related to the postsynapse, such as *Dlg3, Epb41l1, Grin1*, and *Nlgn1*, contribute to the formation of the postsynaptic adhesion molecule complex^29^, called the Neuroligin complex. Neuroligins interact with neurexins, a family of presynaptic adhesion molecules, and are major regulators of synapse development and function^30^. Interestingly, *Inpp5k* has been reported to act as a lysophosphatidic acid signaling modulator, and its overexpression promotes the intrinsic axon growth of corticospinal axons^31^. Considering these reports and the results of our NGS analysis, it is presumed that neurons cultured on the microgrooved nanosheet become more mature than neurons cultured on the flat nanosheet.

To validate the up-regulated DEGs identified by RNA-Seq, we performed quantitative RT-PCR (qRT-PCR) on independently prepared mRNA samples from both flat and microgrooved nanosheets. Since NGS analysis revealed that synapse formation and maturation of the neurons were promoted on microgrooved nanosheets, we selected 14 DEGs known to play crucial roles in either postsynapse or presynapse maturation in neurons: *Ache, Amigo1, Dlg3, Dlgap3, Epb41l1, Epb41l3, Grin, Gripap1, Inpp5k, Kcnd3, Kcnq2, Nlgn1, Pkp4*, and *Stau2*. All the selected DEGs were significantly up-regulated in mRNA samples from neuronal cultures on the microgrooved nanosheets compared with those from the flat nanosheets [*Ache*\*, *Amigo1*\*\*, *Dlg3*\*\*, *Dlgap3*\*, *Epb41l1*\*, *Epb41l3*\*\*, *Grin*\*\*, *Gripap1*\*, *Inpp5k*\*, *Kcnd3*\*\*, *Kcnq2*\*\*, *Nlgn1*\*, *Pkp4*\*\*, and *Stau2\*\** (\*\**p* < 0.01, \**p* < 0.05)] (Figure 4D). These results strongly indicate that microgrooves on a nanosheet can more efficiently facilitate neuronal differentiation, which is consistent with the morphological findings obtained in this study (Figure 2). Importantly, the microgroove structure controls the position of cell adhesion on the nanosheet by limiting it to the bottom and sides of the parallel grooves, which may facilitate neurite bundle formation and lead to the efficient formation of synapses.

In this study, we newly developed a neural cell culture system using a microgrooved PLA nanosheet, which provided a more reproducible and efficient culture environment for the neurons. Thus far, the precise molecular mechanisms by which neuronal maturation is accelerated in the presence of a microgrooved scaffold are still unclear. Nonetheless, this microgrooved nanosheet could provide a powerful means to establish a novel experimental system for neuroscience research and regenerative medicine and may facilitate future investigations of the molecular mechanisms underlying the pathogenesis of many neurological disorders.

## Methods

### Reagents and preparation of nanosheets

All reagents used in this study including the PLA were of analytical grade. Silicon wafers (SiO_2_ substrate; KST World, Fukui, Japan) cut to an appropriate size (typically 3 × 3 cm) were treated with piranha solution, followed by washing with distilled water. PVA (*M*_w_: 22 kDa; Kanto Chemical, Tokyo, Japan) was dissolved in distilled water at a concentration of 10 mg/mL, and this solution was dropped onto the SiO_2_ substrates and spin-coated at 4,000 rpm for 20 s (Spin Coater MS-A100; Mikasa, Tokyo, Japan), followed by drying at 50°C for 2 min. A solution of PLA (*M*_w_: 80–100 kDa; Polysciences, Warrington, PA, USA) at 10 mg/mL was then dropped onto the PVA-coated substrates and spin-coated at 4,000 rpm for 20 s, followed by drying at 50°C for 2 min. The obtained substrates were immersed in distilled water to collect free-standing nanosheets. The nanosheets were scooped up with coverslips and fully dried in a desiccator overnight. PDL or PDL with VTN-N was coated onto the PLA nanosheets immediately prior to use for culture experiments.

### Preparation of microgrooved nanosheets

We prepared PDMS stamps with a microgrooved pattern as previously reported^24,25,26^. The pattern consisted of microgrooves and ridges with a width of 50 μm and a height of 6 μm. To fabricate a free-standing microgrooved nanosheet, we used the same procedures used for a flat PLA nanosheet. These procedures are summarized in Supplementary Fig. 2.

### Coating of nanosheets

For PDL coating, PDL (molecular weight 70–150 kDa, #P6407; Sigma-Aldrich) at a concentration of 0.1 mg/mL in 0.1 M borate buffer was coated onto the nanosheets at a final surface area coating concentration of 30 μg/cm^2^. The nanosheets were then incubated in a 5% CO_2_ incubator at 37°C for 2 h. Next, the nanosheets were washed three times with ultra-pure water and dried on a biological clean bench. For PDL and VTN-N coating, dried PDL-coated nanosheets were incubated with 0.1 mg/mL VTN-N diluted inPBS(–)at 37°C for 2 h in a 5% CO_2_ incubator. The VTN-N solution was removed prior to sticking the Press-to-Seal Silicone Isolator with Adhesive (Thermo Fisher Scientific) onto the nanosheets to control the density of the cells (see “Cell cultures” in METHODS).

### Animals

All animal experimental procedures were approved by The Institutional Animal Care and Use Committee at Tokai University.

### Cell cultures

PC12 cells were cultured in Dulbecco’s Modified Eagle’s medium with High Glucose (Wako) supplemented with 7.5% (w/v) heat-inactivated fetal bovine serum (FBS; PAA Laboratories), 7.5% (w/v) heat-inactivated horse serum (Gibco), 100 U/mL penicillin G, 100 *µ*g/mL streptomycin, and 100 *µ*g/mL sodium pyruvate. Mouse primary cortical neurons were cultured as previously reported.^29,30^ In brief, tissues from each embryo were dissected out and immediately placed into 1 mL of ice-cold HBSS(–). After removing the HBSS(–) by aspiration, 0.5 mL of 0.25% trypsin-EDTA was added and the embryo was incubated for 15 min at 37°C. The trypsin-EDTA was removed, and the embryo was washed several times with 20% FBS/neurobasal medium (Invitrogen). Tissue samples were treated with 50 *µ*g/mL DNase I in 20% FBS/neurobasal medium for 10 min at room temperature (RT). After centrifugation at 150 ×*g* for 15 s, the resulting tissue pellets were dissociated in 0.6 mL of 20% FBS/neurobasal medium by pipetting with a flame-sterilized Pasteur pipette. After counting the number of living cells with the trypan blue assay, 9 × 10^5^ cells were placed onto the PDL- and VTN-N-coated nanosheets using the Press-to-Seal Silicone Isolator with Adhesive to control the cell numbers on the nanosheets. The nanosheets were then immersed in neuronal cell culture media [neurobasal medium containing 1× B-27 supplement (Invitrogen), 25 *µ*g/mL insulin (Sigma-Aldrich), 0.5 mM L-glutamine, 50 *µ*g/mL streptomycin, and 50 U/mL penicillin G] and cultured at 37°C. The medium was then exchanged for fresh medium containing 5% FBS, and the cells were cultured on the nanosheets for another 36 h.

### Cell viability assay

alamarBlue Cell Viability Reagent (Thermo Fisher Scientific) was added to 10% (v/v) of the medium at DIV6 of the primary cultured neurons. After 6 h incubation, we detected the fluorescence intensity of resorufin with the Spectra Max i3 (Perkin Elmer). Resazurin, a PC of the alamarBlue reagent, is reduced to the highly red fluorescent resorufin in viable cells only. Experiments were repeated four times.

### Immunocytochemistry

The cells were fixed with 4% (w/v) paraformaldehyde in PBS(–) pH 7.5 for 30 min at RT and permeabilized with 0.1% (w/v) TritonX-100 in PBS(–) for 30 min. The primary antibodies used in previous reports^32,33^ were diluted in PBS(–) containing 1.5% (v/v) normal goat serum and incubated with the samples. After washing the cells with PBS(–), Alexa 594-conjugated goat anti-mouse IgG (1:500; Molecular Probes) or Alexa 594-conjugated goat anti-rabbit IgG (1:500; Molecular Probes) was used for the detection of proteins of interest. Fluorescent signals were captured with the BZ-X fluorescence microscope (Keyence) and processed with Adobe Photoshop (Adobe).

### Library preparation

Total RNA was extracted from the cultured cells with the RNeasy Plus Micro Kit (Qiagen) according to the manufacturer’s protocol. The quality of the total RNA samples was validated with the RNA 6000 Pico Kit (Agilent) on the Bioanalyzer (Agilent). High-quality RNA samples with an RNA integrity number >9 were used for library preparation. RNA-Seq libraries were prepared with the Encore Complete RNA-Seq DR Multiplex system (NuGEN) in accordance with the manufacturer’s instructions.

### RNA-Seq analysis

Indexed paired-end cDNA sequencing libraries were sequenced by MiSeq (Illumina, San Diego, CA, USA). A total number of 75-bp paired-end reads were sequenced. After trimming the reads with the fastq_quality_trimmer tool in the FASTX-Toolkit (version 0.0.14) using the option (–Q 33, –t 20, –l 30), the reads were then mapped onto the mouse reference genome (mm10) using HISAT2 (version 2.1.0)^34^ with the default options. StringTie (version 1.3.4b)^35^ was used to quantify gene expression. The R package of DESeq2 (version 1.18.1)^27^ was used for RNA-Seq differential expression analysis. We first normalized the gene expression values for each sample using regularized log transformation implemented in the DESeq2 program. For each gene, the gene expression data of cells cultured on both the flat and microgrooved nanosheets were statistically examined. We assumed that genes differentially expressed on these two types of nanosheets with a statistical significance of *p*□<□0.05 (Benjamini–Hochberg corrected) was indeed a DEG. Enrichment analysis was carried out with the enrichGO function in the R package of clusterProfiler (version 3.7.1) ^36^.

### qRT-PCR

qRT-PCR was performed on 10 ng of total RNA using the Thunderbird SYBR qPCR/RT Set (Toyobo) with the specific primers (0.2 *µ*M each) listed in Table 1. The transcript levels were normalized by the amount of *Gapdh* mRNA in each sample. Statistical analyses were conducted with Prism 7 (GraphPad). Statistical significance was evaluated by ANOVA followed by appropriate post hoc tests for multiple comparisons between groups.

**Table 1.**
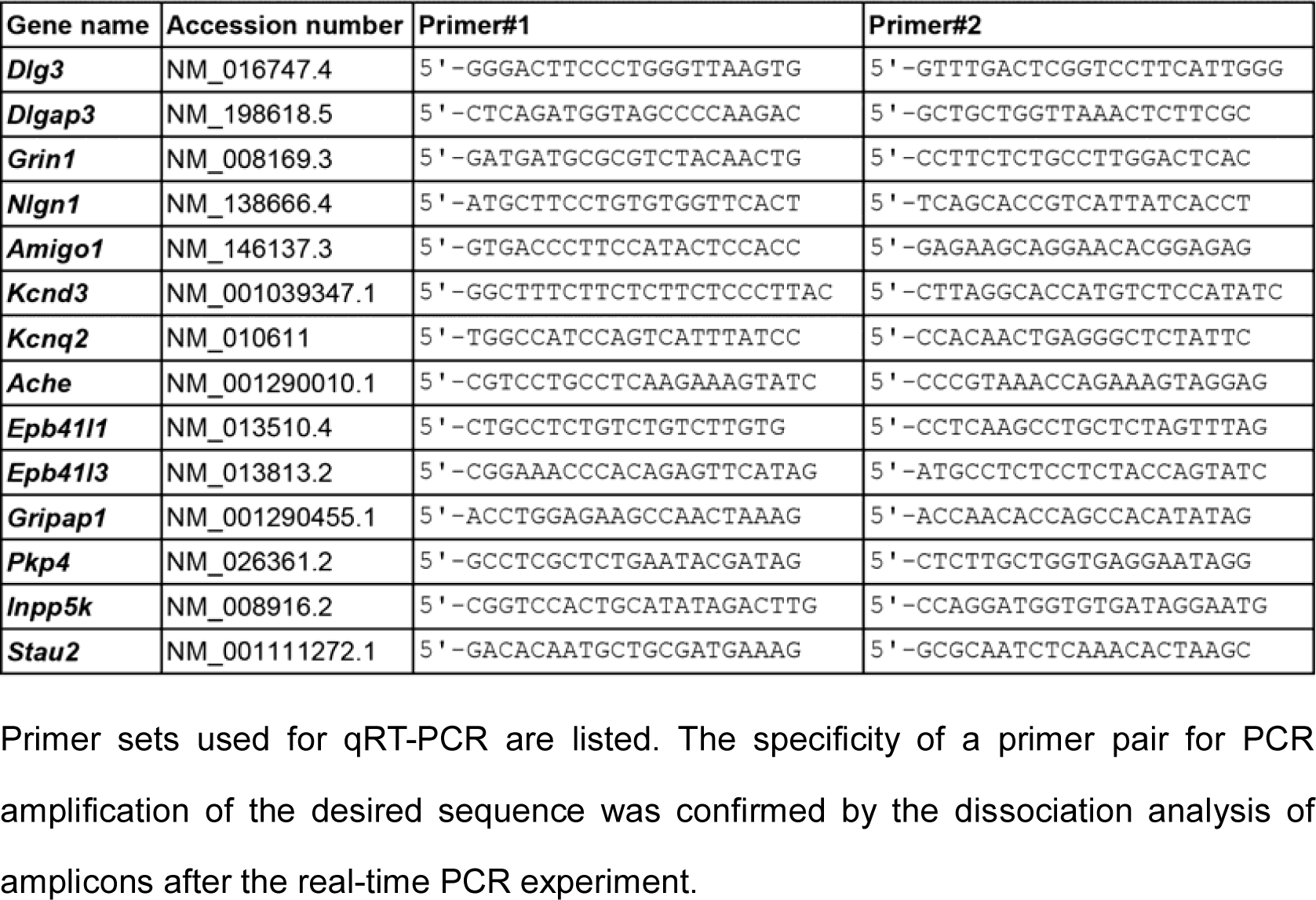
Primer sets used in this study.

## Data Availability

The RNA-Seq data obtained in this study have been deposited in the DDBJ DRA database (https://www.ddbj.nig.ac.jp/dra/index-e.html) under the accession numbers DRR166653–DRR166659.

## Acknowledgments

We thank Hiromi Takahashi and all the members of the Support Center for Medical Research and Education at Tokai University for their technical support in this study. This research was funded by the MEXT (Japanese Ministry of Education, Culture, Sports, Science and Technology)-Supported Program for the Strategic Research Foundation at Private Universities, Grant #S1411010. A.O. was supported by 2015–2016 Tokai University School of Medicine Research Aid. T.F. was supported by the Precursory Research for Embryonic Science and Technology (PRESTO) program from the Japan Science and Technology Agency (JST; grant number JPMJPR152A).

## Author Contributions

A.O., Y.O., and S.N. conceived the study idea. A.O. conducted the biological experiments. M.T.U. and S.N. conducted the data analysis. Y.O., A.H., T.F., and S.T. fabricated the nanosheets. S.H. provided mice for the culture experiments and interpreted the data. A.O., M.T.U., S.H. and S.N. wrote the manuscript. All authors read and approved the final manuscript.

## Additional Information

### Supplementary information

A detailed description of the protocol used to fabricate the microgrooved nanosheet, the results of the surface analysis of the microgrooved nanosheet, and the sequences of the primer sets used in this study are available online.

