## Supporting Information for "Efficient differentiation and polarization of primary cultured neurons on poly(lactic acid) scaffolds with microgrooved structures"

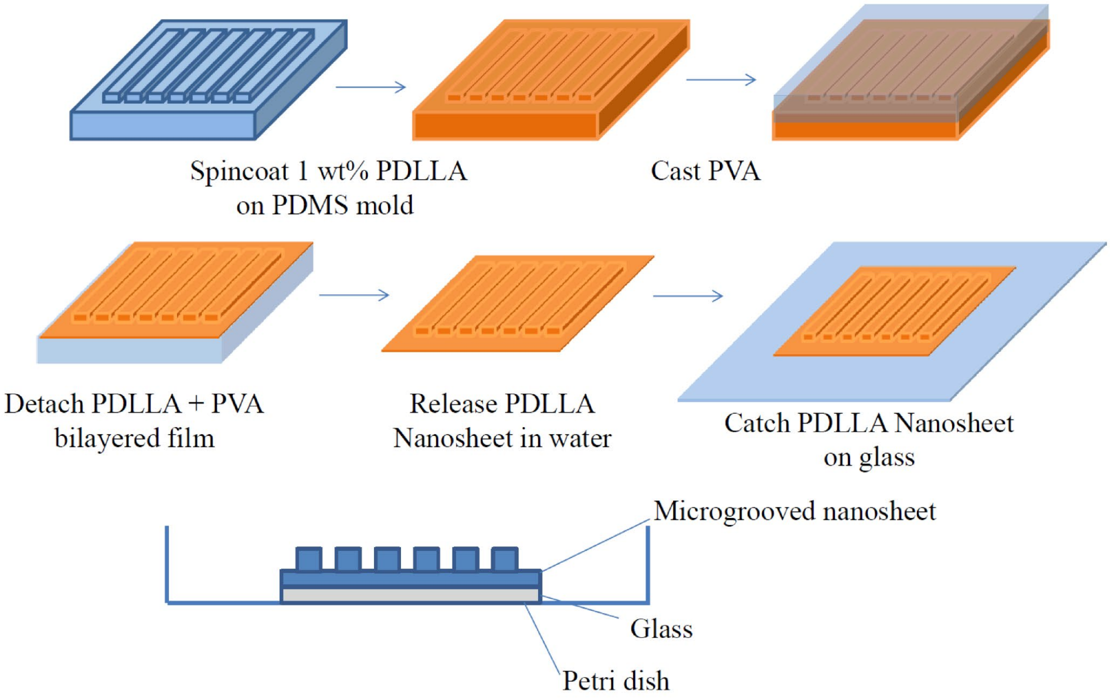


**Figure S1. Protocol for the fabrication of the grooved nanosheets**

The preparation scheme for the microgrooved PLA nanosheets is shown. PLA nanosheets were prepared by spin coating a 10 mg/mL poly(D,L-lactic acid) (PDLLA) solution onto a PDMS negative replica with grooved motifs. Next, a PVA (MW: 13,000–23,000; Kanto Chemical, Inc., Tokyo, Japan) supporting layer was cast onto the PLA nanosheet. The micropatterned PLA nanosheet with the PVA layer was released from the PDMS mold and then placed into a PBS solution to dissolve the PVA supporting layer.

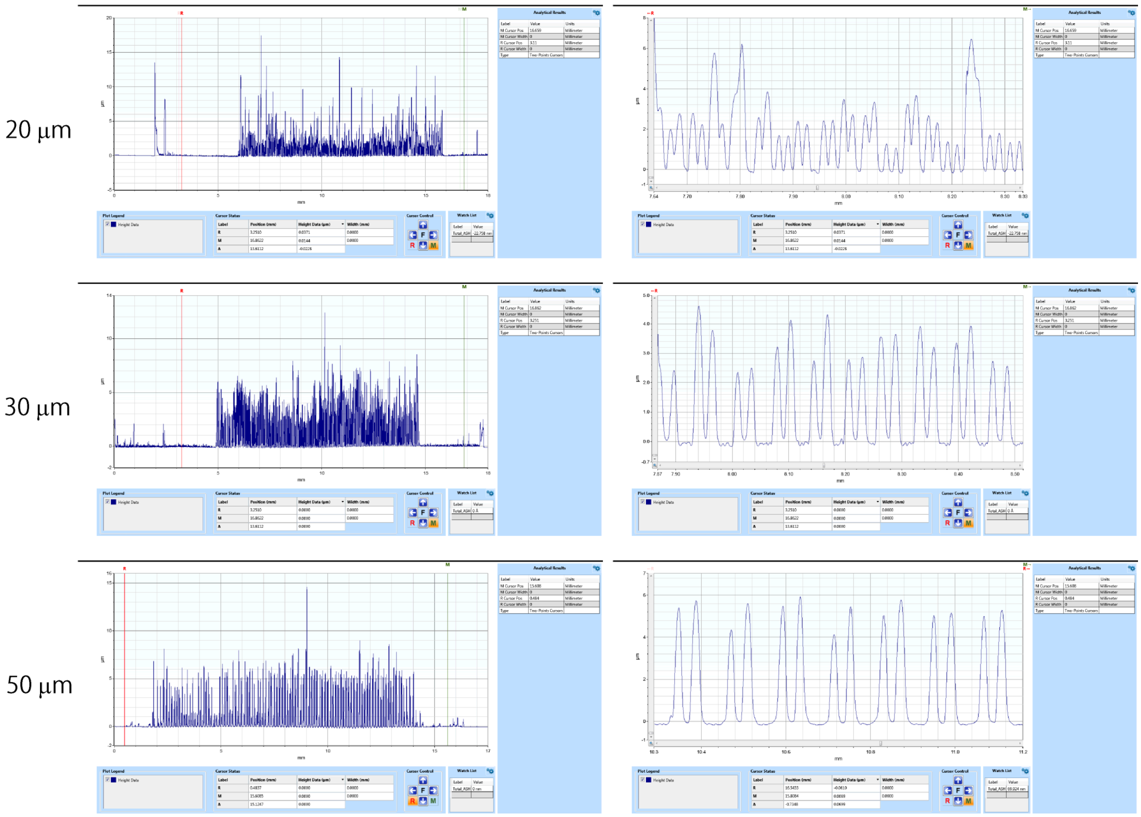


**Figure S2. Surface analysis of the grooved nanosheet**

The thickness and surface morphology of PLA nanosheets with microgrooves of different widths were analyzed with a surface profiler. The microgroove pattern comprising grooves 50 μm in width was the most uniform of the three tested patterns.
